# Coexistence theory and the frequency-dependence of priority effects

**DOI:** 10.1101/243303

**Authors:** Po-Ju Ke, Andrew D. Letten

## Abstract

Priority effects encompass a broad suite of ecological phenomena. Several studies have suggested reframing priority effects around the stabilizing and equalizing concepts of coexistence theory. We show that the only compatible priority effects are those characterized by positive frequency dependence.

## Introduction

The order species arrive in a locality can have lasting impacts on the diversity, composition and function of ecological communities [1, 2]. This phenomenon, alternately referred to as priority effects, alternative/multiple stable states, historical contingency or founder control, was originally explored analytically through Lotka-Volterra competition models [3, 4]. In these simple models, priority effects emerge when competing species have greater negative impacts on heterospecifics than conspecifics, resulting in each species’ growth rate being a positive function of its relative abundance. From a theoretical perspective, the term priority effect is thus strictly defined [5], but over time its usage has broadened to encompass a wider suite of phenomena. Several studies have subsequently mooted the prospect of reorganizing priority effects around the stabilizing and equalizing concepts of coexistence theory [6–8]. Here, we identify the unrecognized problems and promise of such an endeavour.

## The frequency dependence of priority effects

According to coexistence theory, species can coexist when the fitness differences between them are smaller than their niche differences, where the former compares overall adaptedness to a shared environment, and the latter captures overlap in resource usage in space and time [9]. This is equivalent to stating that each species exhibits negative frequency dependence (NFD); i.e., reduced growth as a function of its own relative abundance in a community. For a two-species Lotka-Volterra model, this can be summarized via the inequality 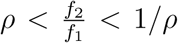, where niche overlap, *ρ*, is equal to ‘1 - the niche difference’ and is bounded between 0 and 1, and 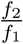 is the fitness ratio. It follows that we can differentiate between two classes of coexistence mechanisms: equalizing mechanisms that reduce the average fitness difference and stabilizing mechanisms that reduce niche overlap.

In addition to being ecologically intuitive, the bounding of niche overlap between 0 and 1 has statistical provenance in Chesson’s original definition as the least squares correlation between the resource utilization functions in MacArthur’s consumer-resource model [10]. More recently, however, Chesson provided a convenient formula for niche overlap in terms of a symmetric measure of the ratio of inter-to intra-specific density dependence based on Lotka-Volterra coefficients, [11]. Specifically, 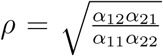. Whether or not a given *ρ* generates NFD depends on the fitness difference between competing species, but it is clear from this formulation that *ρ* is bounded by 0 and 1 only when the product of the intra-specific coefficients is greater than the product of the inter-specific coefficients. When the reverse is true, *ρ* can take values greater than 1.

At first glance, *ρ* > 1 is at odds with both intuitive and statistical interpretations of niche overlap. However, if we redefine niche difference (i.e., 1 - *ρ*) as a measure of stabilization potential, it operationalizes the original analytical definition of priority effects in a form that recognizes the joint role of stabilizing and equalizing mechanisms. More specifically, the criteria for positive frequency dependence (PFD), and therefore stable priority effects, is the inverse of the stable coexistence inequality (Eq. 1), i.e 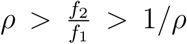, where *ρ* = 1 - stabilization potential [8]. As such, any mechanism that reduces the fitness ratio, or decreases the stabilization potential, will increase the probability of priority effects. Rather than being monotonic, note that the stabilization potential diverges around zero such that values above zero represent the stabilization potential for coexistence, whereas values below zero represent the stabilization potential for priority effects; in other words the strength of the attractor towards alternative stable states. We note that this terminology is different from recent heuristic translations of priority effects into coexistence theory, where niche differences decreasing below zero has been referred to as destabilization [6, 7]. However, in keeping with dynamic systems theory, we favor conceptualizing destabilization as any process that causes the stabilization potential to approach zero from values above or below (Fig 1e in Box 1).

**Figure 1.**
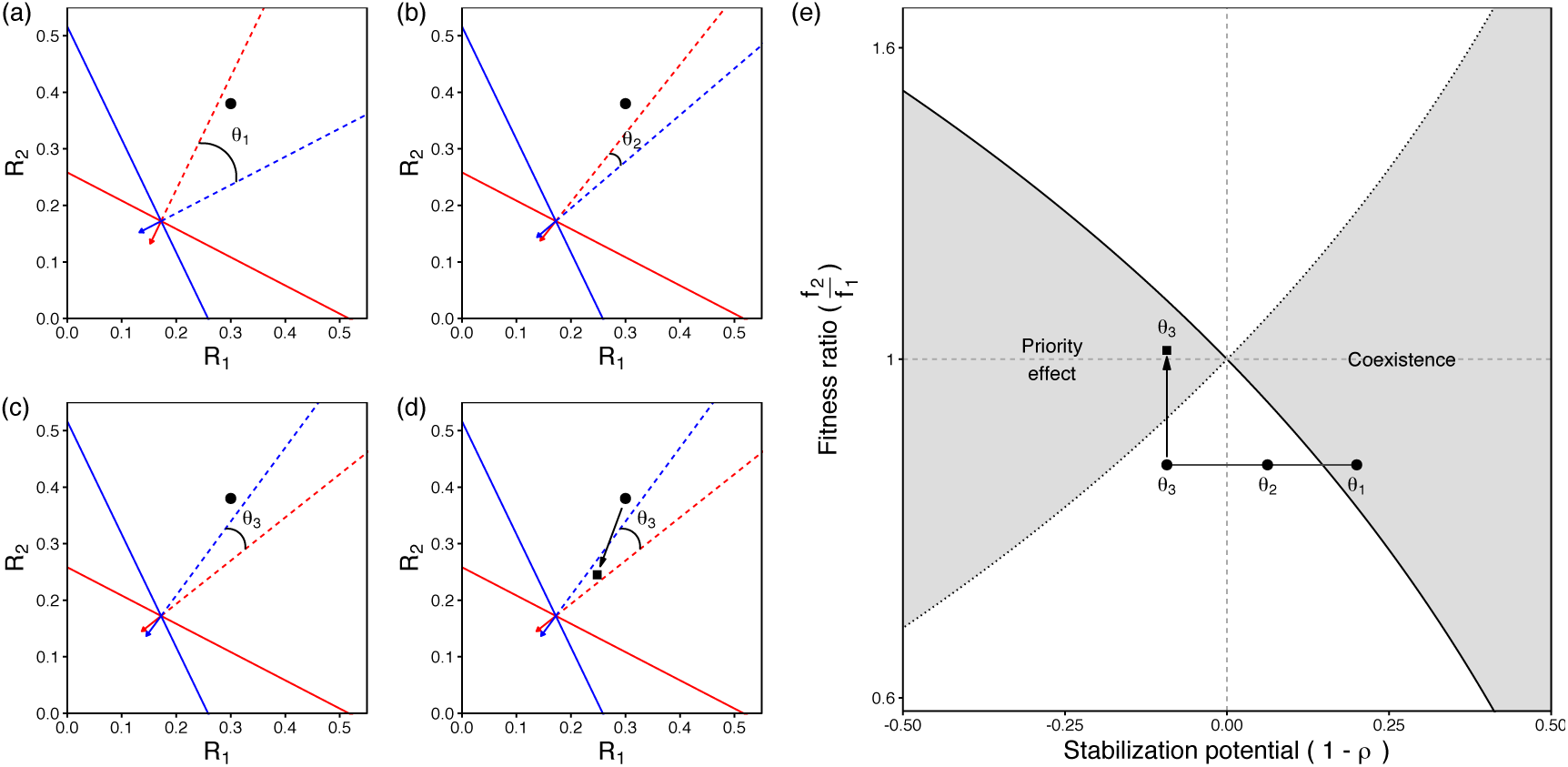
Effect of (a) changing species’ impact vector and resource supply ratio in a consumer-resource model on (e) the fitness ratio and stabilization potential (niche difference) of coexistence theory. In panel (a), the solid red and blue lines are the ZNGIs for each species; the solid lines with arrow heads are the respective impact vectors; the dashed lines are the inverse of the impact vectors; and the black circle and square represent two different supply points that favor blue and red species, respectively. In panel (e), the x-axis represents the stabilization potential (1 - *ρ*) and the y-axis represents the fitness ratio, *f*_2_/*f*_1_; the solid and dotted line represents the boundary where *f*_2_/*f*_1_ equals to *ρ* and 1/*ρ*, respectively; and the right and left gray shaded area indicates the coexistence and priority effect region, respectively. The angles given by *θ*_1–3_ in (a-d) correspond to the respective *θ*_1–3_ in (e). Note that the y-axis is on log-scale.

#### Box 1

A classic example of priority effects emerging from PFD comes from Tilman’s 1982 monograph [12]. Using the approach taken by [8] to derive niche overlap and the fitness ratio from Tilman’s consumer resource model, PFD generated priority effects can be partitioned into stabilizing and equalizing components. In Figure 1a, NFD and coexistence arise due to a combination of intersecting zero net growth isoclines (ZNGIs), consumption vectors directed towards each species’ favored resource, and an intermediate resource ratio. As the angle between the consumption vectors declines to *θ_2_* (Fig. 1b), the stabilization potential also declines. The outcome is competitive exclusion (Fig. 1e). Once the consumption vectors cross and begin to diverge, each species consumes more of its competitor’s favored resource (θ_3_, Fig. 1c), setting up the conditions for PFD. However, if the fitness difference remains sufficiently large, the outcome will still be exclusion irrespective of arrival order (Fig. 1e). If the resource supply shifts to a more balanced ratio (Fig. 1d), the fitness inequality is reduced and priority effects emerge (Fig. 1e). This demonstrates that priority effects are a function of both the stabilization potential and the fitness inequality.

In Box 1 we use Tilman’s consumer-resource model [12] to illustrate that only a subset of phenomena commonly referred to as priority effects are compatible with coexistence theory. In particular, compatible phenomena are limited to those that generate PFD and therefore are consistent with the original definition derived from Lotka-Volterra [5]. This is not to say that PFD is unique to systems exhibiting point equilibria. For example, the coexistence mechanism relative nonlinearity can generate PFD when species that benefit from fluctuations in the intensity of competition also exacerbate those fluctuations [13]. Nevertheless, in a system that precludes the emergence of positive (or negative) frequency dependence, and hence the emergence of a non-trivial stable attractor, the stabilization potential term is unquantifiable. This criterion, however, wholly or partially excludes a number of phenomena, which for heuristic reasons are also routinely termed priority effects. We briefly consider two of these phenomena below.

## Positive density dependence and facilitation

When applying coexistence theory to study priority effects, it is important to recognize that PFD can emerge from negative (*α_ij_ >* 0) or positive (*α_ij_* < 0) density dependence, i.e., facilitation. However, while conceptually compatible with coexistence theory, the analytical tools currently available (i.e., Eq. 3) cannot be leveraged to interpret facilitative dynamics. This is because negative *α_ij_* would generate unbounded population densities unless constrained by specific model designs. As such, Eq. 3 can only be applied to PFD emerging from negative density dependence (e.g. Box 1).

An alternative form of positive density dependence sometimes characterized as a priority effects is an Allee effect [5]. For species exhibiting an Allee effect, there is a density threshold dividing two alternative stable states, i.e., above which the population persists and below which the population goes extinct. As such the alternative stable states arise at the population level, and therefore are distinct from priority effects that emerge at the community level driven by PFD. Thus, while Allee effects can effect community composition if inter-specific interactions maintain species below their Allee threshold, they arise independently of a species’ frequency in a community.

## Succession and transient priority effects

The notion of priority effects has also been usefully applied to understand the effects of arrival order on successional dynamics. In these instances, differences in initial abundance can cause compositional trajectories to vary through time, even though they may all eventually converge on the same community state. In naturally ephemeral microbial systems, such as those that develop in floral nectar or woody debris, this final state might be the local extinction of all community members following the exhaustion of available resources. Such “alternative transient states” [2] that are a outcome of resource pre-emption may have downstream impacts on pollinator preference and decomposition rates, and therefore undoubtedly reflect ecological phenomena with meaningful consequences for ecosystem function. Nevertheless, in the absence of another process sustaining PFD, there is little scope or rationale to understand these phenomena through the lens of coexistence theory.

## Summary

Interest in coexistence theory has been growing steadily, but to date the overwhelming emphasis has been on the underlying stabilizing mechanisms giving rise to NFD and stable coexistence. We have illustrated the most accessible approach to incorporating priority effects mediated through PFD into this body of theory. When priority effects emerge from positive density dependence, or arise in transient systems, it is currently unclear how to analytically connect them to the fundamental concepts of coexistence theory.

## Acknowledgements

We thank Tad Fukami, Tess Grainger, and Daniel Stouffer for comments.

## References

[1] Chase, J.M. (2003) Community assembly: when should history matter? Oecologia 136, 489–498

[2] Fukami, T. (2015) Historical contingency in community assembly: integrating niches, species pools, and priority effects. Annu. Rev. Ecol. Evol. Syst. 46, 1–23

[3] Lewontin, R.C. (1969) The meaning of stability. In Brookhaven symposia in biology, vol. 22. p. 13

[4] May, R.M. (1971) Stability in multispecies community models. Mathematical Biosciences 12, 59–79

[5] Petraitis, P. (2013) Multiple stable states in natural ecosystems. OUP Oxford

[6] Mordecai, E.A. (2011) Pathogen impacts on plant communities: unifying theory, concepts, and empirical work. Ecol. Monograph 81, 429–441

[7] Fukami, T. et al. (2016) A framework for priority effects. Journal of Vegetation Science 27, 655–657

[8] Letten, A.D. et al. (2017) Linking modern coexistence theory and contemporary niche theory. Ecological Monographs 87, 161–177

[9] Chesson, P. (2000) Mechanisms of maintenance of species diversity. Annu. Rev. Ecol. Syst. 31, 343–366

[10] Chesson, P. (1990) MacArthur’s consumer-resource model. Theor. Popul. Biol. 37, 26–38

[11] Chesson, P. (2013) Species Competition and Predation. In R. Leemans, ed., Ecological Systems. Springer New York, pp. 223–256

[12] Tilman, D. (1982) Resource Competition and Community Structure. (Mpb-17). Princeton University Press, Princeton, NJ, USA

[13] Chesson, P. (2009) Scale transition theory with special reference to species coexistence in a variable environment. Journal of Biological Dynamics 3, 149–163

